# FOXO Transcription Factors Activate Alternative Major Immediate Early Promoters to Induce Human Cytomegalovirus Reactivation

**DOI:** 10.1101/2020.02.10.942433

**Authors:** Andrew E Hale, Donna Collins-McMillen, Erik M Lenarcic, Jeremy P Kamil, Felicia Goodrum, Nathaniel J Moorman

## Abstract

A key step during viral reactivation from latency is the re-expression of viral genes. Hematopoietic progenitor cells (HPCs) support human cytomegalovirus (HCMV) latency, and their differentiation triggers cellular cues that drive reactivation. A key step during HCMV reactivation in latently infected HPCs is re-expression of viral genes. We recently determined that the major immediate early promoter (MIEP), which is primarily responsible for MIE gene expression during lytic replication, remains silent during reactivation. Instead, alternative promoters in the MIE locus are induced by reactivation stimuli. Here, we find that forkhead family (FOXO) transcription factors are critical for activation of alternative MIE promoters during HCMV reactivation, as mutating FOXO binding sites in alternative MIE promoters decreased HCMV IE gene expression upon reactivation and significantly decreased the production of infectious virus from latently infected primary CD34^+^ HPCs. These findings establish a mechanistic link by which infected cells sense environmental cues to regulate latency and reactivation, and emphasize the role of contextual activation of alternative MIE promoters as the primary drivers of reactivation.

**Significance:** Human cytomegalovirus infection is lifelong and persistent. Periodic reactivation of cytomegalovirus poses serious disease risk for immune-compromised patients. A critical driver of reactivation is expression of viral genes from the major immediate early locus. Recent paradigm-shifting evidence shows that reactivation is driven from promoters distinct from those that drive replication in permissive cells. Understanding the contextual control of these promoters and how they specifically respond to cellular cues that drive reactivation is critical for establishing future therapies that prevent reactivation. Our findings mechanistically define a previously enigmatic relationship between differentiation and reactivation, and provide potential targets for therapeutic intervention to prevent HCMV reactivation and disease.

## Introduction

Reactivation of latent human cytomegalovirus (HCMV) infection poses a significant threat to patients with compromised immune systems. While primary infections in healthy adults are typically associated with little or mild disease, reactivation in immunocompromised individuals, such as solid organ and stem cell transplant recipients and can lead to significant morbidity and mortality(1). Despite this, little is known regarding the mechanisms controlling HCMV reactivation.

Following an initial burst of viral gene expression upon entering cells that support latency (e.g. CD34^+^ hematopoietic stem cells; HPCs), HCMV gene expression is largely silenced, allowing the virus to persist in a quiescent or latent state(2, 3). Pro-inflammatory cytokines and cellular cues that drive differentiation induce the re-expression of viral lytic cycle genes, culminating in the production of infectious virus that can spread throughout the host and cause disease(4, 5). Critical to HCMV reactivation is the re-expression of the viral immediate early 1 and 2 proteins (IE1 and IE2), which drive the expression of the HCMV lytic cycle(6-8). Defining factors that regulate IE1 and IE2 expression during reactivation is thus critical for understanding the mechanisms controlling reactivation and resulting HCMV disease.

In cells permissive for lytic replication such as fibroblasts, the expression of both IE1 and IE2 transcripts is largely driven by the major immediate early promoter (MIEP), which is silenced during latency. Until recently it was presumed that reactivation of the lytic cycle relied on the resumption of MIEP activity. However we recently discovered two alternative promoters (iP1 and iP2, together referred to as intronic promoters) within the first intron of the canonical major immediate early locus(9). Rather than inducing MIEP activity, HCMV reactivation stimuli instead induce transcription from the iP1 and iP2 promoters, which correlates with the increase in IE1 and IE2 mRNA levels(10). Deletion of the intronic promoters significantly attenuates the production of infectious virus after reactivation, revealing that the iP1 and iP2 promoters play critical roles in IE1 and IE2 re-expression and HCMV reactivation.

Here we begin to unravel the mechanisms by which HCMV senses cellular cues to trigger IE1 and IE2 re-expression by defining FOXO transcription factors as key players linking cellular differentiation to HCMV reactivation. These data provide new mechanistic insights into HCMV reactivation and suggest that manipulating FOXO transcription factor activity may be a future means to limit HCMV disease.

## Materials and Methods

### Cells and Viruses

MRC-5 fibroblasts and HeLa cells were grown in Dulbecco’s modified Eagle medium (DMEM) supplemented with 10% fetal bovine serum and penicillin/streptomycin. HCMV TB40E derived from a bacterial artificial chromosome (BAC) containing an SV40 promoter-driven GFP reporter (11) was used as wild type virus, and served as the backbone for the generation of recombinant viruses. Titers of cell free virus were determined by the 50% tissue culture infective dose (TCID50) method on MRC-5 fibroblasts. Unless otherwise noted, all infections were performed at a multiplicity of infection (MOI) of 1 for one hour, followed by removal of the inoculum.

### Construction of Recombinant Viruses

BAC-mediated recombineering was used to generate viral mutants on the TB40E genomic background using a two-step recombination approach, as before (9). Briefly, the iP2 locus was replaced with a kanamycin/levansucrase fusion cassette (KanSacB) using homology mediated recombination in recombination competent SW105 E.coli (12). The KanSacB cassette was then replaced with the iP2 sequences containing the indicated mutation in a second round of recombination. The entire genomes of the wild type and recombinant BACs were then sequenced to ensure no additional changes were present other than the intended mutations.

### THP-1 Latency Model

Latency studies were conducted essentially as described previously (10, 13, 14). Briefly, cells were with HCMV (TB40/E; (5, 15)) at a multiplicity of 2. After 5 days cells were treated with 100 nM 12-o-<u>t</u>etradecanoyl<u>p</u>horbol-13-<u>a</u>cetate (TPA) to induce reactivation, or with DMSO solvent control. Whole cell lysates, DNA, and RNA were collected at each of the indicated time points.

### Assay of Infectious centers for latency and reactivation

The frequency of HCMV reactivation in CD34^+^ HPCs was quantified as previously described (10, 14). Briefly, pure populations of CD34^+^ HPCs were infected with TB40E WT or FOXO3mut123 recombinant virus (MOI = 2). Twenty-four hours after infection, latently infected cells were purified and co-cultured with stromal cells to maintain latent infection (10, 14). After 10 days, latently infected cells were co-cultured with naïve MRC-5 fibroblast monolayers to quantify the number of infectious centers produced.

### Quantitative real-time PCR analysis (qRT-PCR)

mRNA abundance was quantified by qRT-PCR essentially as described previously (10, 16). Briefly, RNA was extracted with TRIzol and then treated with DNase. The RNA was reverse transcribed and quantified, and then quantified by real-time PCR using SYBR green incorporation and transcript-specific primers. For transfected cells, RNA abundance was determined by comparison to a standard curve generated from qPCR analysis of 10-fold serial dilutions of a DNA standard specific for each primer pair. Transcript abundance in latently infected cells was measured as previously described (10). Detailed methods are provided in the Supplementary Materials and Methods.

### Plasmid construction

Luciferase reporter vectors containing the dP, MIEP, iP1, or iP2 promoters have been previously described (9). Details of additional plasmids used in this study are described in the Supplemental Materials and Methods.

### Luciferase assays

Luciferase assays were performed as described previously (17). Luciferase activity was measured 24 hours after transfection of Hela cells, and normalized to the protein concentration in the sample. Luciferase reactions were performed in duplicate for each sample and averaged, and the graphs show the mean values of at least three biological replicates performed on different days.

### Electrophoretic mobility shift assays (EMSA)

Double stranded DNA probes were incubated with 500ng purified recombinant FOXO3a protein (Abnova) on ice for 20 minutes. The reaction was resolved on polyacrylamide gel and then transferred to a nylon membrane. Membranes were probed with streptavidin-HRP according to manufacturer’s instructions (ThermoFisher) and visualized by chemiluminescence (BioExpress).

### Western Blotting

Western blotting was performed as described previously (18). Briefly, cells were lysed in RIPA buffer containing protease inhibitors and equal amounts of protein were resolved on SDS-PAGE gels. Proteins were transferred to nitrocellulose membranes (Amersham) and probed with primary antibodies followed by horseradish peroxidase-coupled secondary antibodies. Membranes were visualized by chemiluminescence (ECL) reagent (BioExpress) using a chemiluminescence detection system (Bio-Rad).

### In silico analysis of FOXO binding sites in the MIE locus

The program Find Individual Motif Occurrences (FIMO;(19)) was used to search intron A of the MIE genomic locus for the two known FOXO binding motifs RWAAAYAA and MMAAAYAA. Only sites with a threshold p value of p<.01 were considered.

## Results

We recently described a novel mechanism by which the re-expression of the IE1 and IE2 mRNAs during HCMV reactivation is driven by the alternative promoters iP1 and iP2 (Fig 1A), rather than the MIEP (10). While iP1 and iP2 promoters are necessary for efficient reactivation, the factors regulating their activity are unknown. Reactivation is induced by differentiation of monocytes into macrophages, a process that is mimicked in experimental latency models by treating latently infected cells with phorbol esters (e.g., TPA) (10) or LY294002 (20), a chemical inhibitor of the PI3K/mTOR pathway. A literature review found that phorbol esters, LY294002, and monocyte differentiation all activate FOXO family of transcription factors (21). An *in silico* analysis of the HCMV MIE locus identified three potential FOXO binding sites upstream of iP2 (Fig 1B), the most active MIE promoter during reactivation (10). We thus hypothesized that FOXO transcription factors stimulate iP1 and iP2, resulting in increased transcription of the IE1 and IE2 mRNAs and HCMV reactivation.

**Figure 1.**
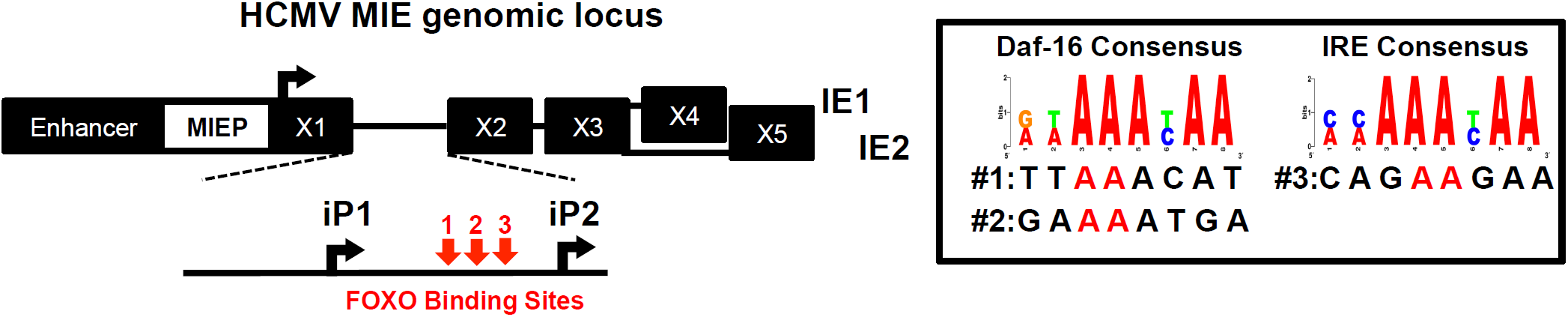
HCMV intronic promoters contain FOXO TF consensus sites. (A) Schematic showing the location of the intronic promoters iP1 and iP2 and potential FOXO binding sites in the the major immediate early intronic promoter locus. (B) The intronic promoters were searched against the two consensus binding motifs for FOXO transcription factors. Three sequences of the three potential FOXO binding sites are shown. The bases highlighted in red were changed to XX to inn the mutants described in Figures 2-6.

To test this hypothesis, we first determined if FOXO transcription factors (TFs) stimulated the activity of the iP1 and iP2 promoters outside the context of infection. We co-transfected vectors expressing FOXO1 and FOXO3a with reporter constructs containing the distal promoter (dP), MIEP, iP1, or iP2 promoters upstream of luciferase. We found neither FOXO1 nor FOXO3a affected the activity of the MIEP or the dP, but both FOXO1 or FOXO3a significantly increased the activity of both iP1 and iP2 (Fig 2A,B). We mutated critical nucleotides in each potential FOXO binding site in iP2, the most FOXO-responsive promoter, (shown in Fig 1B), and also generated a reporter construct containing mutations in all three binding sites to determine if FOXO binding to multiple sites had a cooperative effect. Mutating any of the three potential FOXO binding sites, either alone or in combination, significantly decreased iP2 induction by FOXO transcription factors (Fig 2 C, D).

**Figure 2.**
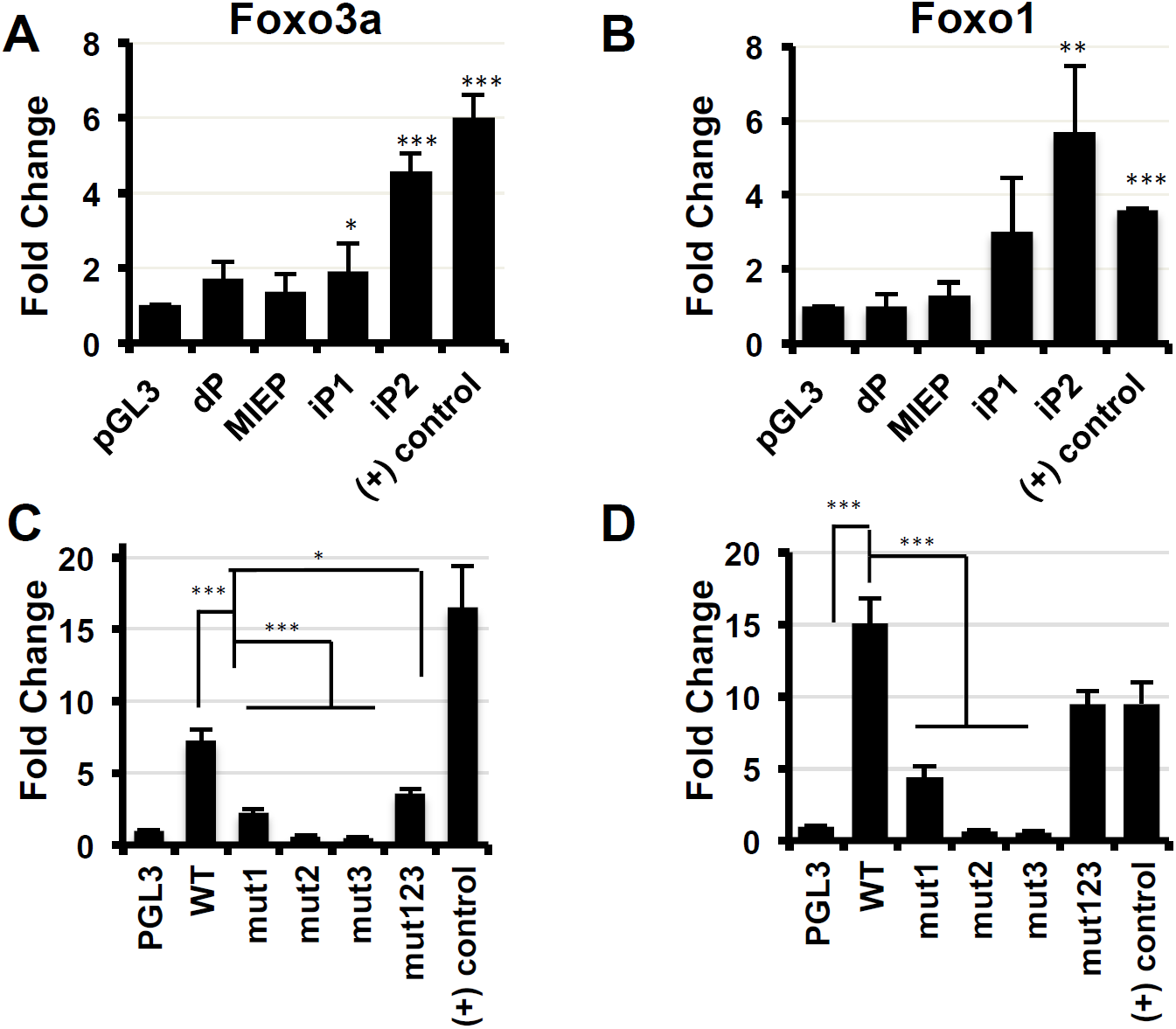
A) MIE Intronic promoters are activated by FOXO. (A and B) Hela cells were co-transfected with FOXO overexpression plasmids and previously described luciferase reporter constructs containing the four promoters in the MIE genomic locus 5’ of the luciferase gene; the distal promoter (dP), the major immediate early promoter (MIEP) intronic promoter 1 (iP1) and intronic promoter 2 (iP2). Luciferase activity was measured at 24 hours after transfection and normalized to the amount of protein in the sample. A previously characterized positive control ((+) control) was included which contains three consensus FOXO binding sites 5’ of the luciferase gene. The parental luciferase plasmid pGL3 Basic, which lacks a promoter upstream of luciferase, served as a control. The graphs show the fold change in each promoters activity in the presence of FOXO3a (A) or FOXO1 (B) compared to the pGL3 Basic control. (C and D) Each potential FOXO binding site in the iP2 promoter was mutated either alone (mut1, mut2, mut3) or in combination (mut123).The reporters were co-transfected with either FOXO3a (C) or FOXO1 (D), and luciferase assays were performed as above. (n=3; * = *p* < 0.05, ** = *p* < 0.05, *** = *p* < 0.005)

We next determined how FOXO transcription factors affect IE1 and IE2 expression in the context of the MIE locus. To this end, we measured IE1 and IE2 protein levels in cells co-transfected with FOXO expression vectors and the plasmid pSVH, which contains the entire MIE locus, from 840 base pairs upstream of the MIEP transcription start site (TSS) through the 3’UTR of exon 5 (22-24). In this construct, the MIEP is the predominant promoter driving IE1 and IE2 transcription, and both iP1 and iP2 are minimally active (9). We found expression of FOXO3a, but not FOXO1, resulted in a slight, but reproducible, increase in IE1 and IE2 protein levels (Fig S1).

To confirm that FOXO TFs increase IE1 and IE2 expression independently of MIEP, we repeated the above experiments using a variant of the pSVH plasmid where the core MIEP is deleted (pSVHΔMIEP). We previously showed that IE1 and IE2 expression from this plasmid is driven exclusively by the iP1 and iP2 promoters (9). As before, deleting the core MIEP promoter greatly decreased IE1 and IE2 protein levels (Fig. 3A). In contrast to the luciferase reporters, FOXO1 expression had minimal affect on IE1 or IE2 protein levels in this setting. However, FOXO3a expression significantly increased IE1 and IE2 protein levels (Fig 3A). Surprisingly IE1 and IE2 protein levels were comparable to levels in cells transfected with the wild type MIE locus in the presence of FOXO3a (Fig. 3A).

**Figure 3.**
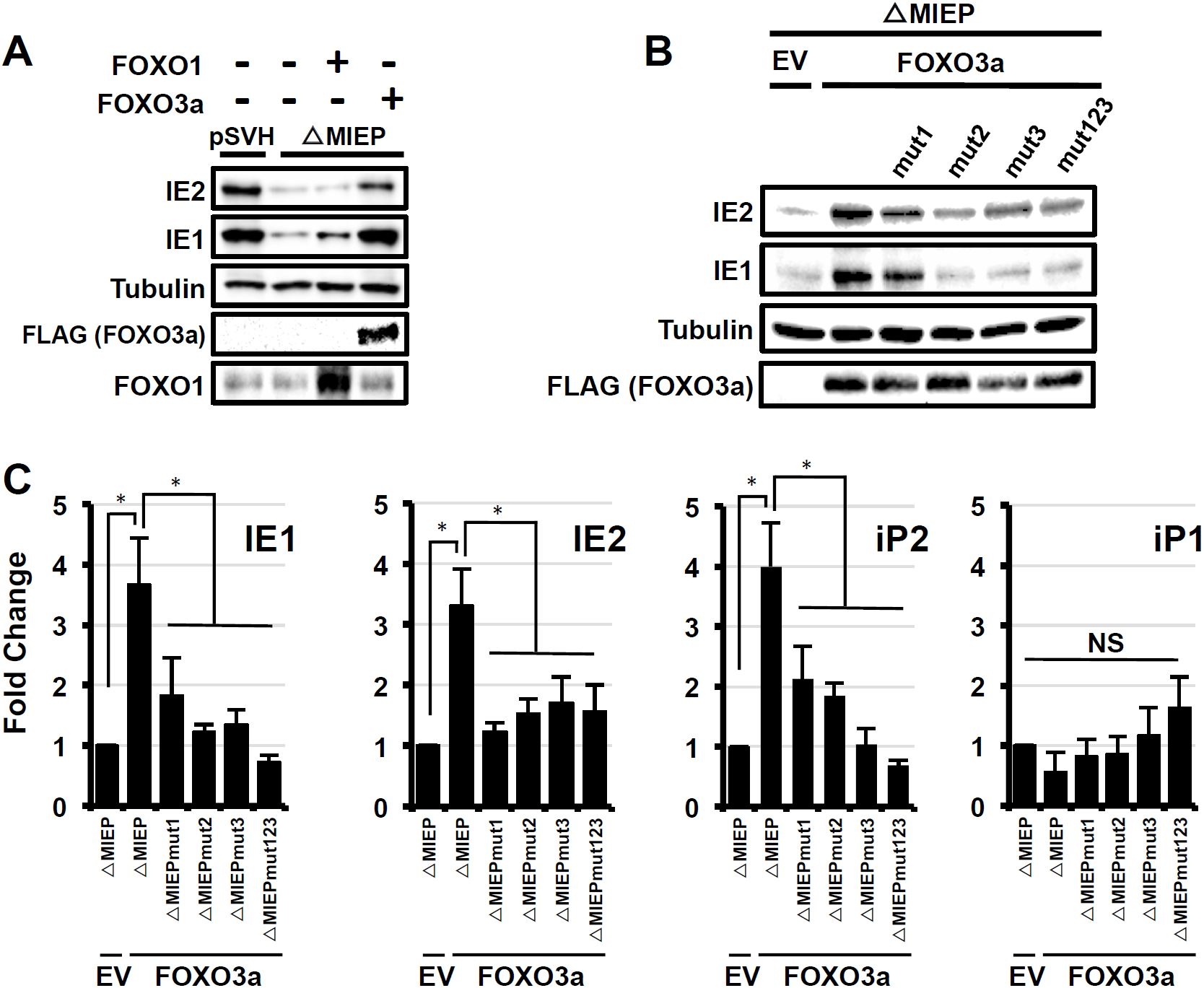
FOXO3a stimulates transcription from iP2 in the context of the MIE genomic locus. (A) A plasmid containing the regulatory and coding regions for the HCMV IE1 and IE2 genes (pSVH) or a variant of pSVH lacking the core MIEP (pSVHΔMIEP) was transfected into Hela cells together with a FOXO1 or FOXO3a expression plasmid, or an empty expression vector (EV) control. Cells were harvested at 24 hours after infection, and cell lysates were analyzed by Western blot using the indicated antibodies. Results are representative of three independent experiments. (B) Each FOXO consensus site in iP2 was mutated in the context of the pSVHΔMIEP plasmid, either alone (mut1, mut2, mut3) or in combination (mut123). Cells were transfected with the indicated plasmid along with a FOXO3a expression plasmid (FOXO3a) or the empty vector control (EV) and harvested 24 hours later. Cell lysates were analyzed by Western blot using the indicated antibodies. The results are representative of three independent experiments. (C) Cells were transfected and harvested as in B. RNA was extracted and analyzed by qRT-PCR using primers specific for IE1, IE2, iP1, or iP2. The graphs show the fold change in RNA abundance compared to cells transfected with pSVHΔMIEP and the empty expression vector control. (n=3; * = *p* <0.05)

As FOXO1 had minimal impact on IE1/2 expression in the context of the MIE locus, we focused further studies on FOXO3a. Mutating any of the potential FOXO TF binding sites abrogated the effects of FOXO3a on IE1 and IE2 protein (Fig 3B) and mRNA levels (Fig 3C). The increase in IE1 and IE2 mRNA correlated with a matching increase in transcription from the iP1 and iP2 promoters (Fig 3C), suggesting that FOXO3a increases IE1 and IE2 expression by stimulating iP1 and iP2 activity.

To determine if FOXO TFs directly bind FOXO-responsive sites in the MIE intronic promoters, we used an electrophoretic mobility shift assay (EMSA) to measure binding of recombinant FOXO3a to double stranded oligonucleotides containing each of the three potential FOXO binding sites. FOXO3a bound to each of the three sites, with FOXO Site 3 exhibiting the greatest binding (Fig 1 & 4A). Mutating each of the potential FOXO sites using the same strategy as above decreased FOXO3a binding to each site (Fig A,B). Together with the results above, these data show FOXO3a can directly bind specific sequences within the iP2 promoter, suggesting that the effect of FOXO3a on IE1 and IE2 expression is not due to an indirect effect on other cellular pathways.

**Figure 4.**
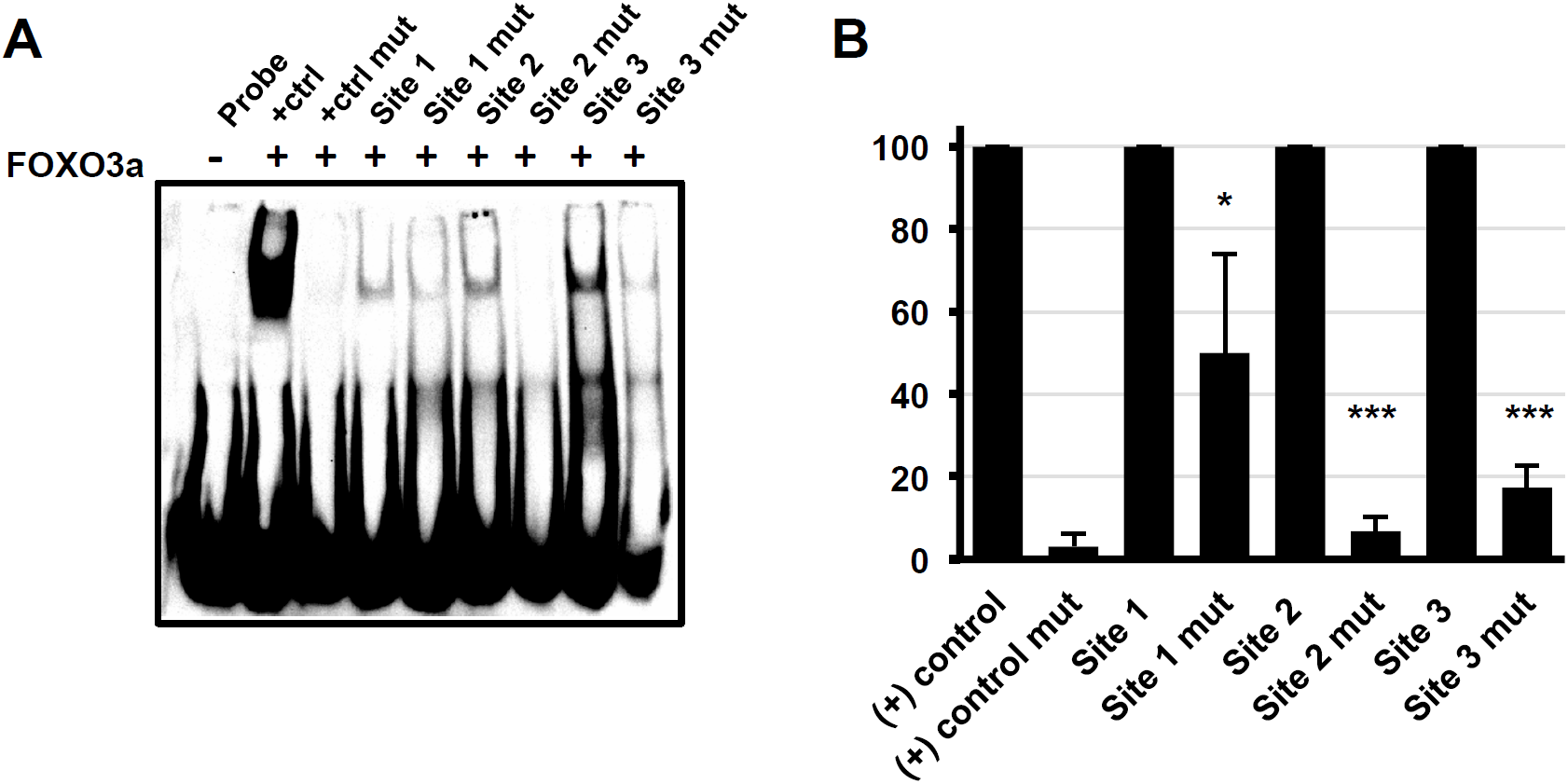
FOXO3a directly binds to FOXO consensus sites in iP2. (A) Recombinant purified FOXO3a was incubated with double stranded biotinylated probes containing the indicated FOXO consensus sites in iP2 (Site 1, Site 2, Site3), or mutated sequences containing the changes shown in Figure 1 (Site 1 mut, Site 2 mut, Site 3 mut). Protein:DNA complexes were resolved on non-denaturing acrylamide gels, and the complexes were visualized by chemiluminescence. The data are representative of three independent experiments. (B) Graph showing the decrease in FOXO3a binding to the mutant probes compared to the wild type probes, which are set to 100%. The graph shows the mean of three independent experiments. (* = *p* < 0.05; ** = *p* < 0.005; *** = *p* <0.0005

We next sought to define the role of FOXO TF stimulation of the iPs in HCMV reactivation. We considered depleting FOXO TFs from latently infected cells, however FOXO TFs play critical roles in cellular differentiation (25). Therefore any reactivation phenotypes found in FOXO-depleted, latently infected cells could be due to defects in differentiation, defects in intronic promoter activation, or both. To circumvent this issue, we generated a recombinant virus containing mutations in the FOXO binding sites in iP2 and measured the effects on virus replication and latency. In fibroblasts the recombinant virus expressed viral immediate early (IE), early (E), and late (L) proteins (Fig. S2B) similarly to the wild type virus control, and replicated to equivalent titer and with similar kinetics as wild type virus in fibroblasts (Fig. S2A). These results are consistent with our previous results showing a minor role for iP1 and iP2 in HCMV lytic replication in fibroblasts (9, 10).

To determine the role of FOXO binding to the iP1 and iP2 promoters in regulating IE1 and IE2 expression in hematopoietic cells, we infected THP-1 cells, a model system for HCMV latency studies, with wild type virus or the recombinant containing mutations in all three FOXO binding sites. THP-1 cells are an established model of HCMV latency and reactivation, which we previously used to demonstrate a requirement for iP1 and iP2 in reactivation (Fig 5A) The homogeneity of THP-1 cells offers the advantage of a synchronous establishment of latency and re-expression of MIE genes following a reactivation stimulus (TPA) that is not possible with primary hematopoietic cells. THP-1 cells were infected to similar levels with WT and mutant virus, as determined by the percentage of GFP-positive cells (Fig 5B). Similar to infection with viruses lacking the entire iP1 and iP2 elements (10), mutating FOXO binding sites in iP2 decreased IE protein accumulation immediately after infection and prior to the establishment of latency. In cells allowed to establish a quiescent infection, the recombinant virus also expressed significantly less IE1 and IE2 protein (Fig 5C) after the addition of TPA, a reactivation stimulus. As previously shown, transcripts derived from the MIEP remained very low both before and after TPA treatment of cells infected with wild type virus, and importantly MIEP-derived transcript levels were not significantly affected by the absence of the FOXO binding sites. In contrast, iP1 and iP2 transcripts accumulated to high levels in cells infected with wild type virus before being silenced for latency, and were induced following TPA stimulation (Fig 5D). iP1 and iP2 transcripts were expressed to significantly lower levels in cells infected with the FOXO binding site mutant relative to the wild type virus over the time course, and were re-expressed to lower levels following TPA treatment. From these data we conclude that FOXO binding sites in the intronic promoters play an important role in the initial early burst of MIE expression, and TPA-induced MIE re-expression in THP-1 cells.

**Figure 5.**
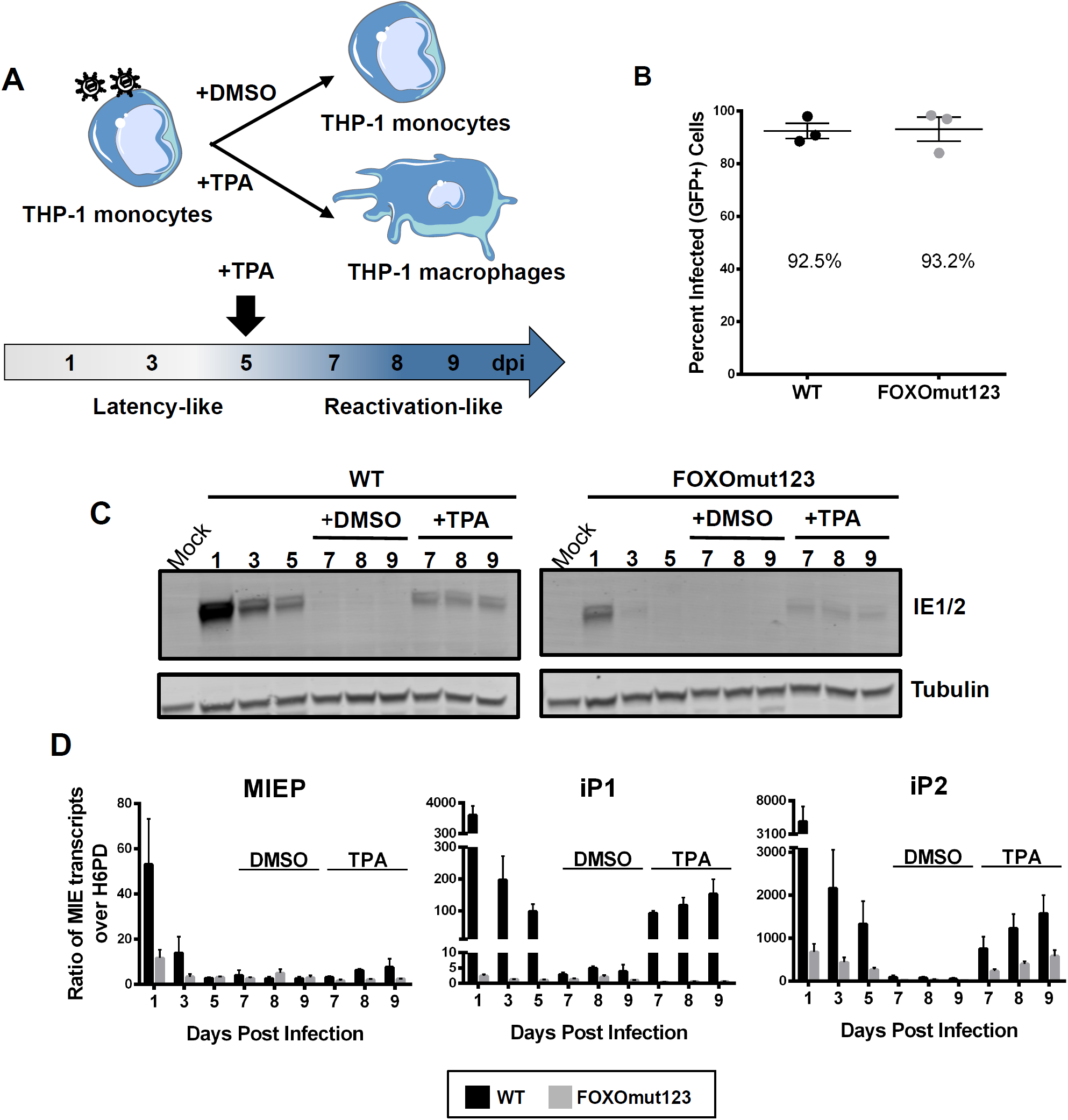
FOXO consensus sites in IP2 are required for re-expression of MIE genes in THP-1 cells. (A) Schematic of THP-1 model for re-expression of MIE genes after stimulation with TPA. THP-1 cells were infected with TB40/E WT or FOXOmut123. At 5 dpi, cells were treated with TPA to stimulate macrophage differentiation and re-expression of viral genes, or with a DMSO vehicle control. (B) Equal infection (GFP+) of THP-1 cells with each virus was determined by flow cytometric analysis at 24 hpi. Biological replicates are represented as single points around the mean. Standard error is shown. (C) Cellular lysates were harvested at the indicated time points and immunoblotted to detect HCMV IE1 and IE2 and tubulin, as a loading control. (D) MIE transcripts derived from the MIEP, iP1, or iP2 were quantified using qRT-PCR relative to the low-abundance housekeeping gene H6PD. Error bars represent the average of three independent experiments, each in triplicate. The standard error of the mean is shown. Significant differences in the number of iP1 and iP2 transcripts (but not MIEP) were found at 1, 3, and 5 dpi and 7, 8, and 9 dpi with TPA (*p* ≤0.05).

We also compared wild type and FOXO binding site mutant virus reactivation in primary CD34^+^ bone marrow-derived HPCs, the gold standard model for HCMV latency studies (26-28). Pure populations of infected CD34+ HPCs were isolated and cultured for 10 days. The frequency of infectious centers were quantified following reactivation or in a lysate prepared prior to reactivation. We observed a significant decrease in the amount of infectious virus produced following reactivation of the recombinant virus as compared to wild type virus (Fig 6). Together these data show that while dispensable for lytic replication, FOXO sites in the intronic MIE promoters play critical roles in HCMV reactivation.

**Figure 6.**
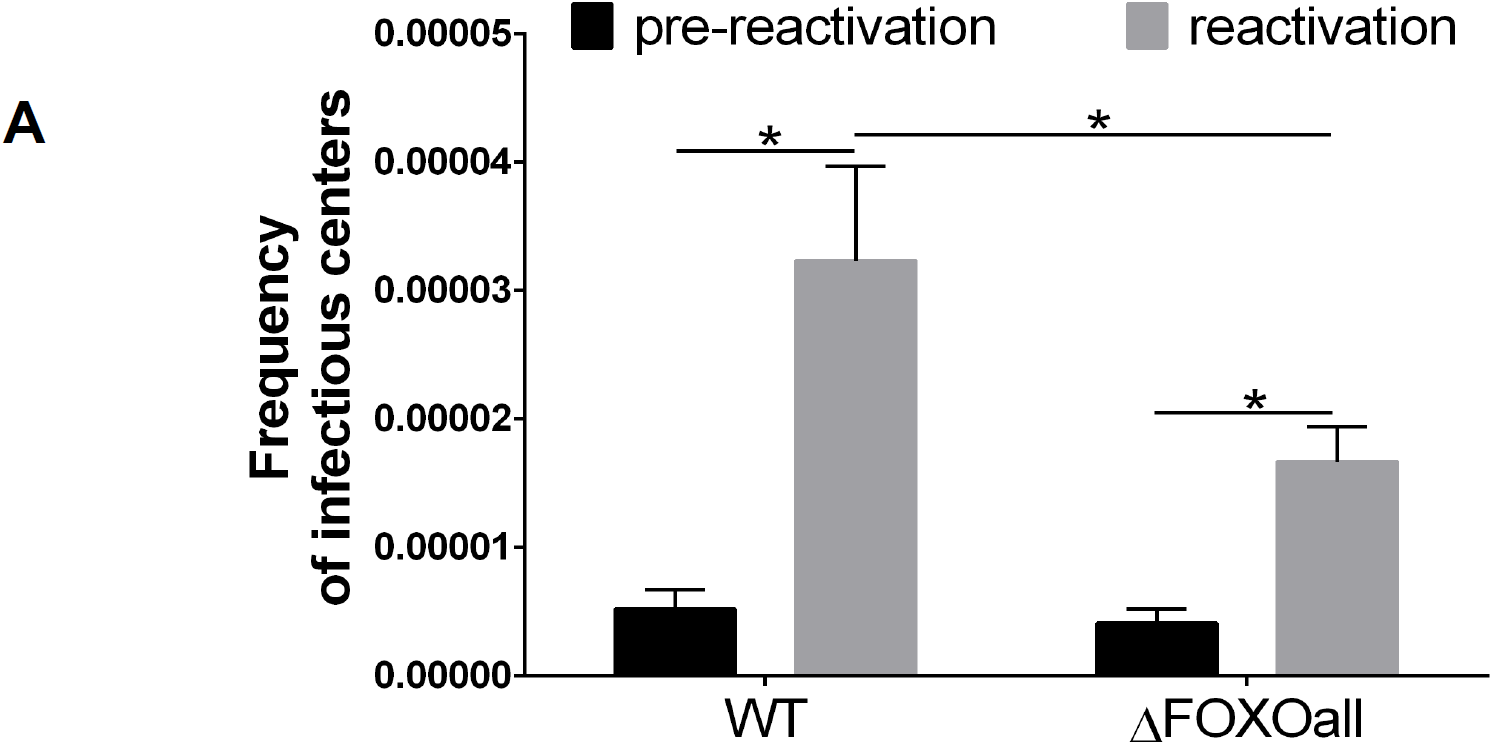
FOXO binding sites in iP2 are necessary for efficient HCMV reactivation in CD34+ HPCs. CD34+ HPCs were infected with TB40/E WT or the mutant viruses where all three FOXO binding sites were mutated (FOXOmut123) at an MOI of 2. Pure populations of infected (GFP+) CD34+ HPCs were isolated by FACS and seeded into long term bone marrow cultures. At 10dpi, viable CD34+ HPCs or an equivalent cell lysate (pre-reactivation control) were seeded by limiting dilution onto fibroblast monolayers in a cytokine-rich media to promote myeloid differentiation. Infectious centers were scored 14 days later and expressed as the frequency of infectious centers. Bars represent standard error of the mean. Statistical significance was determined using 2-way analysis of variance (ANOVA) with repeated measures by both factors (wild-type vs. mutant and pre-reactivation vs. reactivation where * indicates *p* ≤0.05).

## Discussion

Contrary to the long-held paradigm that viral MIE gene re-expression during reactivation requires the MIEP, we recently discovered two novel promoters in the MIE locus, iP1 and iP2 (9), that are necessary for IE gene re-expression and HCMV reactivation (10). Here we begin to unravel the regulatory mechanisms controlling iP1 and iP2 activity by showing that the FOXO transcription factors are critical positive regulators of MIE gene expression from iP1 and iP2 in hematopoietic cells. As FOXO transcription factors also drive cellular differentiation, our results provide a mechanism by which HCMV senses and responds to changes in cellular differentiation to regulate reactivation.

Both FOXO1 and FOXO3a can activate the intronic promoters, however our results suggest that FOXO3a is the critical FOXO transcription factor required for reactivation. While FOXO1 increased intronic promoter activity from a luciferase reporter, FOXO1 did not significantly impact IE1 and IE2 expression in the context of the more complex MIE genomic locus. While FOXO1 and FOXO3a are both expressed in hematopoietic cells, FOXO3a more efficiently localizes to the nucleus in myeloid progenitor cells, particularly during cellular stresses such as those that induce reactivation(29, 30). Further, FOXO3a regulates monocyte to macrophage differentiation (25, 31), while FOXO1 promotes maintenance of stemness in CD34^+^ HPCs (25). Further, FOXO3a directly binds specific sequences in the intronic promoters (Fig 4) and strongly increases their activity (Fig 2). While additional studies are needed to more fully elucidate the role of specific FOXO TFs in HCMV biology, these data suggest that FOXO3a is a particularly crucial positive regulator of viral lytic gene expression in the contexts of HCMV latency and reactivation.

Our data also provide new insight into the roles of intronic promoters in the context of latency and reactivation. While deleting either iP1 or iP2 decreases reactivation, transcripts arising from the iP2 promoter were more abundant upon reactivation (10). We find that FOXO TFs are critical for IE1 and IE2 expression in hematopoietic cell infection, which correlates with the FOXO-dependent accumulation of both iP1 and iP2 transcripts (Fig 5). The effect of FOXO binding site mutation on both iP1 and iP2 activity further suggests that transcription from these elements is coordinated. Importantly, our data show that FOXO binding sites in iP1 and iP2 are critical for the early burst of MIE gene expression immediately after infection of HPCs, consistent with the requirement for iP1 and iP2 for efficient MIE gene expression early after infection of HPCs (10). Our data also show that FOXO binding sites in the intronic promoters are critical for efficient reactivation (Fig 6), suggesting that activation of iP1 and iP2 by FOXO TFs is also critical for MIE gene expression during reactivation. Going forward, it will be important to understand how this early burst of MIE gene expression impacts latent infection and reactivation, and how FOXO TFs regulate MIE expression during different stages of infection.

While we define a role for FOXO TFs in reactivation, our data also suggest additional factors regulate intronic promoter activity and HCMV reactivation, as mutation of all three FOXO binding sites in iP2 did not result in a complete loss of reactivation, highlighting the complexity of the MIE locus. Several transcription factors are implicated in reactivation (7, 8, 32-34) suggesting they may also regulate intronic promoter activity. For example, AP1 transcription factor binding sites in the MIE enhancer are required for efficient IE gene re-expression and intronic promoter activation during reactivation (Christine O’Connor, personal communication). Further studies are needed to identify additional transcription factors that regulate the intronic promoters in conjunction with FOXO TFs to control HCMV reactivation.

## Acknowledgements

We wish to thank Jim Alwine, John Purdy, Stan Lemon, Mark Heise, Ralph Baric, Blossom Damania and members of the Moorman, Goodrum and Kamil labs for helpful conversations and critical feedback. This work was supported by NIH grants AI143191 to NJM, JP and FG, AI127335 to FG, AI123811 and AI103311 to NJM, and support from the UNC Virology Training Grant (T32 AI07419) to AEH. DCM is supported by a Postdoctoral Fellowship (18POST33960140) from the American Heart Association.

