## Supplemental Figures for "FOXO Transcription Factors Activate Alternative Major Immediate Early Promoters to Induce Human Cytomegalovirus Reactivation"

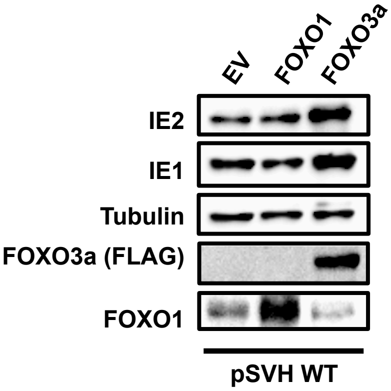


**Supplemental Figure 1. FOXO3a increases IE1 and IE2 expression from the MIE genomic locus.** A) Western blots of Hela cells co-transfected as in Figure 3 with pSVH and FOXO expression vectors or the empty expression vector (EV) control. The results are representative of at least three independent experiments.

### Supplemental Figure 1

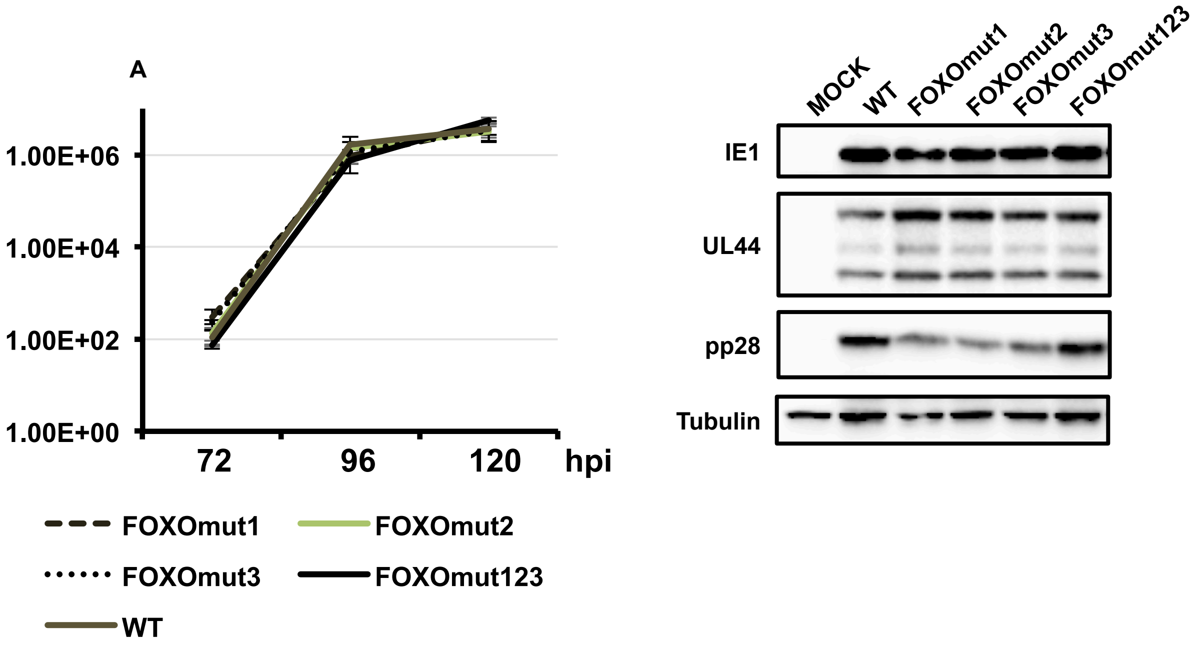


**Supplemental Figure 2. FOXO binding sites in iP2 are dispensable for HCMV lytic replication. (**A) Titer of wild type (WT) or FOXO binding site mutants (FOXOmut1, FOXOmut2, FOXOmjut3, FOXOmut123) at 120 hours post infection (hpi) as determined by TCID50 assay on MRC-5 fibroblasts. (B) Immediate early (IE1), early (UL44) and late (pp28) gene expression of FOXO binding mutant viruses at 120 hours post infection (hpi). All experiments were performed three times on three separate days. Representative images from a single experiment are shown.

### Supplemental Figure 2
