## Supplemental Materials and Methods for "FOXO Transcription Factors Activate Alternative Major Immediate Early Promoters to Induce Human Cytomegalovirus Reactivation"

**Construction of Recombinant Viruses**. BAC-mediated recombineering was used to generate viral mutants on the TB40E/GFP genomic background (1, 2) using a two-step recombination approach, as before (3). In the first step, a kanamycin/levansucrase fusion cassette (KanSacB) was PCR amplified with primers containing 50-nucleotide homology arms flanking the targeted insertion site. The PCR product was transformed into recombination competent SW105 E.coli as before (4), and plated on LB agar containing chloramphenicol and kanamycin. The KanSacB cassette was removed by a second round of recombineering using synthetic DNA fragments (see Table S1) and the recombination strategy described above. Following transformation cells were plated on LB agar containing chloramphenicol and 6% sucrose to select against colonies that retained the Kan/SacB cassette, and then tested for kanamycin sensitivity to ensure loss of the kanamycin cassette. The absence of genomic rearrangements was confirmed by restriction digestion of the recombinant BAC DNAs at each recombination step. No unintended mutations were detected when the 500 bp on either side of each mutation were sequenced, or when the entire genome of the recombinant was sequenced. Next-generation sequencing libraries were prepared from BAC DNA using the Nextera library preparation kit (Illumina), and the libraries were sequenced on an Illumina MiSeq instrument. Recombinant BAC DNA was purified using the Nucleobond Midiprep kit (Macherey-Nagel) according to the manufacturer's directions. Infectious virus was reconstituted by transfecting recombinant DAC DNA (5µg) together with pCGN pp71 (1µg; (5) into MRC-5 fibroblasts.

**THP-1 Latency Model.** Latency studies were conducted essentially as described previously (6-8). THP-1 cells were cultured in Roswell Park Memorial Institute (RPMI) medium supplemented with 10% FBS, 1 mM sodium pyruvate, 2 mM glutamax, 50 µM β-mercaptoethanol, and 100 U/mL pen/strep. Cells were infected in suspension ($5e^5$ cells per mL) with TB40E HCMV at an MOI of 2 (as determined by $TCID_{50}$ using MRC-5 fibroblasts). Infected cell suspension was mixed by rocking every 30 minutes for 4 hours, followed by a centrifugal enhancement at 450 x *g* for 20 minutes, yielding 50 – 80% infection ($GFP^+$ cells by flow cytometry) at 24 hours following infection. Infected cells were cultured for 5 days post infection in non-tissue culture-treated cell culture flasks and continued to proliferate; RPMI was added to maintain a cell density of less than $1e^6$ cells per mL. On day 5, cells from each experimental group were pooled and centrifuged at 120 x g for 7 minutes, then resuspended at 5e$^5$ cells per mL. Cells were treated with 100 nM 12-o-tetradecanoylphorbol-13-acetate (TPA) and plated on tissue culture-treated plates to promote monocyte-to-macrophage differentiation (and viral reactivation) or treated with DMSO solvent control and cultured in non-tissue culture-treated flasks to continue the latent infection. Cells were washed in PBS at 24 hours post TPA/DMSO treatment and fresh media was added to the culture at 1 mL for every 5e$^5$ cells for non-adherent cells or the volume required to cover the plate for adherent cells. Whole cell lysates, DNA, and RNA were collected at each of the indicated time points.

**Assay of Infectious centers for latency and reactivation.** The frequency of HCMV reactivation in CD34$^+$ HPCs was quantified as previously described (6, 8). The frequency of HCMV reactivation in CD34$^+$ HPCs was quantified as previously described (6, 8). Briefly, pure populations of CD34$^+$ HPCs were infected with TB40E WT or FOXO3mut123 recombinant virus (MOI = 2) expressing GFP as a marker for infection. At 24 hpi, infected cells (GFP$^+$/CD34$^+$) were isolated by fluorescence activated cell sorting (FACS) and cultured over stromal cells to maintain latent infection. At 10 dpi, viable CD34$^+$ HPCs were seeded by limiting dilution onto monolayers of permissive MRC-5 fibroblasts (reactivation). An equivalent number of CD34$^+$ HPCs were mechanically lysed and seeded in parallel to quantify infectious virus present prior to reactivation (pre-reactivation). Frequency of infectious centers formed prior to and following reactivation was quantified 14 days later by extreme limiting dilution analysis of GFP$^+$ wells.

**Quantitative real-time PCR analysis (qRT-PCR).** mRNA abundance was quantified by qRT-PCR essentially as described previously (6, 9). Total RNA was extracted using TRIzol according to the manufacturer's directions and as described previously (9). Briefly, cells were resuspended in TRIzol, extracted with chloroform, and RNA precipitated with isopropanol. The RNA was treated with Turbo DNase (Applied Biosystems) and quantified on a NanoDrop spectrophotometer. For quantitative reverse transcriptase PCR (qRT-PCR), cDNA was generated from 0.5 µg total RNA using the High Capacity cDNA reverse transcription kit (ThermoFisher) using random hexamers. The abundance of each transcript was determined using a real-time PCR machine (Bio-Rad) and SYBR Select master mix (ThermoFisher). The abundance of each product was determined by comparison to a standard curve generated from qPCR analysis of 10-fold serial dilutions of a DNA standard specific for each primer pair. For comparative analysis of MIE transcript abundance, each of the promoter-specific primer pairs were shown previously to only amplify the intended promoter specific sequence, have similar real-time PCR efficiencies, and have a similar linear range of detection as compared to the appropriate DNA standard (3). Table S1 lists primers used in this study.

For latently infected THP-1 cells, we used a qRT-PCR approach designed specifically to quantify lower abundance transcripts, as before (6). $5e^5$ cells were collected by centrifugation (120 x *g*, 7 minutes) and resuspended in Zymo *Quick*-DNA/RNA Miniprep lysis buffer. Adherent cells were washed with PBS then scraped in DNA/RNA lysis buffer. Total RNA was extracted using the Zymo Duet *Quick*-DNA/RNA miniprep kit (Zymo Research). cDNA was synthesized using the Transcriptor First Strand cDNA Synthesis Kit (Roche). Briefly, total RNA (400ng) was combined with 2.5 µM Anchored-oligo(dT)18 primers and denatured at 65°C for 10 minutes. A Reverse Transcriptase (RT) master mix (1x Transcriptor RT Reaction Buffer, 40 U/µL Protector RNase Inhibitor, 10 mM Deoxynucleotide Mix, 20 U/µL Transcriptor RT) was added to the template-primer mix (RT- control was made by substituting water for RT in a single reaction). Samples were incubated in a Mastercycler (Eppendorf) for 60 minutes at 50°C, then at 85°C for 5 minutes to inactivate the reverse transcriptase. Final reaction products were diluted 1:4 in PCR water to reduce $MgCl_2$ concentration then amplified in a quantitative polymerase chain reaction (qPCR) using 1x Lightcycler 480 SYBR Green I Master Mix (Roche) and 0.5 µM of a series of sequence-specific primer pairs (see Table S1 for detailed target sequences). Each qPCR reaction was run in triplicate; the average cycle threshold was used to quantify viral mRNAs. Relative expression of each mRNA was calculated using the Pfaffl method that accounts for efficiency of individual primer pairs. Primer efficiencies were calculated using an internal standard curve made with cDNA from lytically infected fibroblasts.

**Plasmid construction** Luciferase reporter vectors containing the dP, MIEP, iP1, or iP2 promoters have been previously described (3). To reporters containing mutations in FOXO binding sites, synthetic 500 nucleotide sequences with mutations in the FOXO binding sites (Table S1) were amplified by PCR and cloned into the pGL3-Basic vector (Promega) cut with HindIII and NotI using Gibson assembly (NEB). The same mutations were incorporated into the plasmid pSVH, which contains the entirety of MIE locus from the enhancer region upstream of the MIE promoter to beyond the 3' UTR of exon 5 (10), or a modified version of pSVH where the core region of the MIEP was deleted (pSVHΔMIEP;(3)). Briefly, the synthetic DNAs containing FOXO site mutations were PCR amplified. Either pSVH or pSVHΔMIEP was also PCR amplified, and the PCR product was cloned into pSVHΔMIEP using the Gibson assembly method (NEB). A list of primers used is found in Table S1.

**Luciferase assays** Luciferase assays were performed as described previously (11). Briefly, $1.5e^5$ HeLa cells were seeded in a 12-well tissue culture plate and transfected with the indicated plasmids using polyethylenimine (PEI). Twenty four hours after transfection, the cells were lysed in 150 μl of passive lysis buffer (Promega). Eight microliters of lysate was mixed with 40 μl of luciferase reagent (Promega), and luciferase was measured on a luminometer (Molecular Devices). The luciferase readings were normalized to total protein in each sample as determined by the Bradford protein assay kit (VWR). Luciferase reactions were performed in duplicate for each sample and averaged, and the graphs show the mean values of at least three biological replicates performed on different days. Plasmids used for expression of indicated proteins were: FLAG-Foxo3a WT (Addgene #8360), FLAG-Foxo1 (Addgene #13507), Forkhead Responsive Element positive control, FHRE-Luc (Addgene #1789), pEGFP (Clontech), and pcDNA3.1 (Invitrogen).

**Electrophoretic mobility shift assays (EMSA).** Double stranded biotinylated DNA probes (IDT) were annealed to their reverse complement by briefly heating to 95°C and allowing to slowly cool to room temperature. Double stranded probes were then gel purified using a 15% native PAGE gel containing 1% TBE. Purified probe concentration was determined using a Qubit fluorometer (Invitrogen). 20ng of each probe was incubated on ice with 500ng purified recombinant FOXO3a protein (Abnova, H00002309-P01) in binding buffer (12mM HEPES pH 7.9, 4mM Tris pH 7.9, 60mM KCl, 1mM DTT, 0.6 mg/mL BSA, 10% glycerol and 0.5ug poly DI:DC) for 20 minutes. FOXO3a binding was resolved using a 4% native PAGE gel containing 0.5% TBE followed by semi-dry transfer to nylon membrane (Thermo), UV crosslinked, and probed with streptavidin-HRP according to manufacturer's instructions. Probes sequences are listed in Table S1.

**Western Blotting.** Western blotting was performed as described previously (12). Briefly, cells were lysed in RIPA buffer (50 mM Tris pH 7.4, 150 mM NaCl, 1 mM EDTA, 1% NP-40, 1% sodium deoxycholate) with cOmplete protease inhibitors (Roche) on ice for 10 min and insoluble material was removed by centrifugation. Protein concentration was measured by the Bradford assay, and equal amounts of protein (30 μg) were loaded in each lane. Sample buffer (45 mM Tris pH 6.8, 30% glycerol, 2% sodium dodecyl sulfate, 150 mM DTT, and 180 μM bromophenol blue) was added to each sample prior to boiling at 95°C for 10 min. The samples were then resolved on SDS-polyacrylamide gels (PAGE). Proteins were transferred to nitrocellulose membranes (Amersham) and blocked in Tris-buffered saline with Tween 20 (TBST; 20 mM Tris pH 7.6, 140 mM NaCl, 0.1% Tween 20) with 5% nonfat milk for at least 1 hour at room temperature. For monoclonal antibodies, the membranes were incubated with the indicated dilution of antibody in TBST with 1% bovine serum albumin (BSA) for 2 hours at room temperature or overnight at 4°C. For polyclonal antibodies, the membranes were incubated with the antibody in TBST with 5% BSA overnight at 4°C. Following incubation with primary antibody, the membranes were washed three times with TBST and then incubated with horseradish peroxidase-coupled secondary antibodies for 1 hour at room temperature in TBST with 1% BSA. To visualize resolved proteins, the membranes were incubated with WesternBright enhanced chemiluminescence (ECL) reagent (BioExpress) and imaged on a chemiluminescence detection system (Bio-Rad). The following antibodies were used at the indicated dilution: M2-FLAG (1:1,000, Sigma), FOXO1 (1:1,000, CST), UL44 (ICP36; 1:1,000; Virusys), tubulin (1:10,000; Sigma), GFP (1:2,000; Roche). The IE1 (1:100; (13)), IE2 (1:100; (14)), and pp28 (1:100; (15)) antibodies were generous gifts from Tom Shenk.
